# Neuron types and distribution of a Drosophila driver line collection

**DOI:** 10.64898/2026.10.06.756975

**Authors:** Lisa Mais, Claire Managan, Brianna Yarbrough, Gary Patrick Hopkins, Jennifer Rivas Salinas, Shirley A. Lauchie, Peter Hirsch, Christoph Karg, Stephan Saalfeld, Wyatt Korff, Gudrun Ihrke, Dagmar Kainmueller, Geoffrey W. Meissner

**Author notes:** **For correspondence:** (GWM); (LM). These authors jointly supervised this work (Dagmar Kainmueller, Geoffrey W. Meissner).

## Abstract

Cell type designations label uniquely-identifiable groups of related cells that typically function in similar ways. Cell types divide the *Drosophila melanogaster* central nervous system into functional units, but remain challenging to target and identify. Previously we generated a driver line and image resource for targeting many neuron types (***Meissner et al., 2025***). Here we improve on identifying the targeted neurons by automated searching against electron microscopy datasets and manual scoring of search results to add 6,368 annotations across 1,772 driver lines in the collection. These annotations provide a potential training resource for future search algorithms. We then estimate the number of cell types and thus the fraction of the central nervous system labeled by the collection as a whole.

## Introduction

Neuroscience studies in *Drosophila melanogaster* and related species are made more powerful and informative through the targeting of specific neural populations. The GAL4/UAS and split-GAL4 systems are common approaches to achieve this targeting (***Fischer et al., 1988; Brand and Perrimon, 1993; Jenett et al., 2012; Luan et al., 2006; Tirian and Dickson, 2017; Dionne et al., 2018***). Uniquely identifying the labeled neurons enables comparisons across studies and datasets, but can be challenging.

Morphological search algorithms enable the comparison of neurons identified via electron microscopy (EM) or light microscopy (LM) as long as the volumes have been aligned to a common template, such as JRC2018 (***Bogovic et al., 2020***). NBLAST enables comparison between fully segmented neurons, which are routinely generated for EM datasets, but are more challenging to obtain from LM datasets, which typically contain many more biological samples with limited resolution and higher background levels (***Costa et al., 2016***). Color depth MIP (Maximum intensity projection) search operates efficiently on 2D brain images with depth encoded by color and partial neuron segmentation (***Otsuna et al., 2018; Meissner et al., 2023; Clements et al., 2024***). PatchPerPix neuron segmentation and PatchPerPixMatch (PPPM) search build on NBLAST to search complex or fragmented neuronal expression patterns (***Mais et al., 2020, 2021***). Automation makes the comparison between hundreds of thousands of neuron representations feasible, but match quality remains below the level of trained observers.

The exceptional resolution, connectivity information, complete and monolithic nature, and intense study of EM volumes has recently made them the standard for *Drosophila* cell type information based on anatomy (***Scheffer et al., 2020; Takemura et al., 2024; Marin et al., 2024; Cheong et al., 2026; Azevedo et al., 2024; Schlegel et al., 2024; Nern et al., 2025; Berg et al., 2025***). Comparisons between volumes can help address the idiosyncrasies of individual organisms, proofreading efforts, and nomenclatures (***Schlegel et al., 2024; Nern et al., 2025; Berg et al., 2025; Stürner et al., 2025***). EM/LM cell type matching helps guide EM neuron identification based on LM data, and driver lines characterized in LM are needed for physiological and behavioral studies even when guided by EM connectomes. Thus, mapping EM-derived cell types onto LM data is critical for enabling the effective use of driver line collections and accelerating the identification of relevant lines for downstream biological studies.

To this end, we present an addendum to the recently published split-GAL4 study (***Meissner et al., 2025***). We applied an automated EM/LM matching approach followed by manual proofreading, resulting in the annotation of ∼3,000 Confident cell type-to-line associations across the brain and ventral nerve cord (VNC; Figure 1). These curated associations provide additional EM-derived cell type information for the split-GAL4 collection and serve as a valuable resource for future studies, including the development and evaluation of improved EM/LM matching methods. Leveraging these annotations, we revisited the question of how comprehensively the split-GAL4 collection captures the cellular diversity of the fly central nervous system (CNS), which we refer to as its *coverage*. Although ***Meissner et al. (2025***) estimated that approximately 90% of voxels are covered by the split-GAL4 collection, this spatial measure does not directly quantify the fraction of neuronal cell types represented in the collection. We therefore employed Monte Carlo simulations, parameterized from the empirical cell type annotations, to account for the non-uniform, package-structured sampling of cell types across driver lines (i.e., multiple cell types per line). Based on this analysis, we estimate that approximately one-quarter to one-half of brain and VNC cell types are represented in the collection, providing an alternative metric to the previous voxel-based estimate.

**Figure 1.**
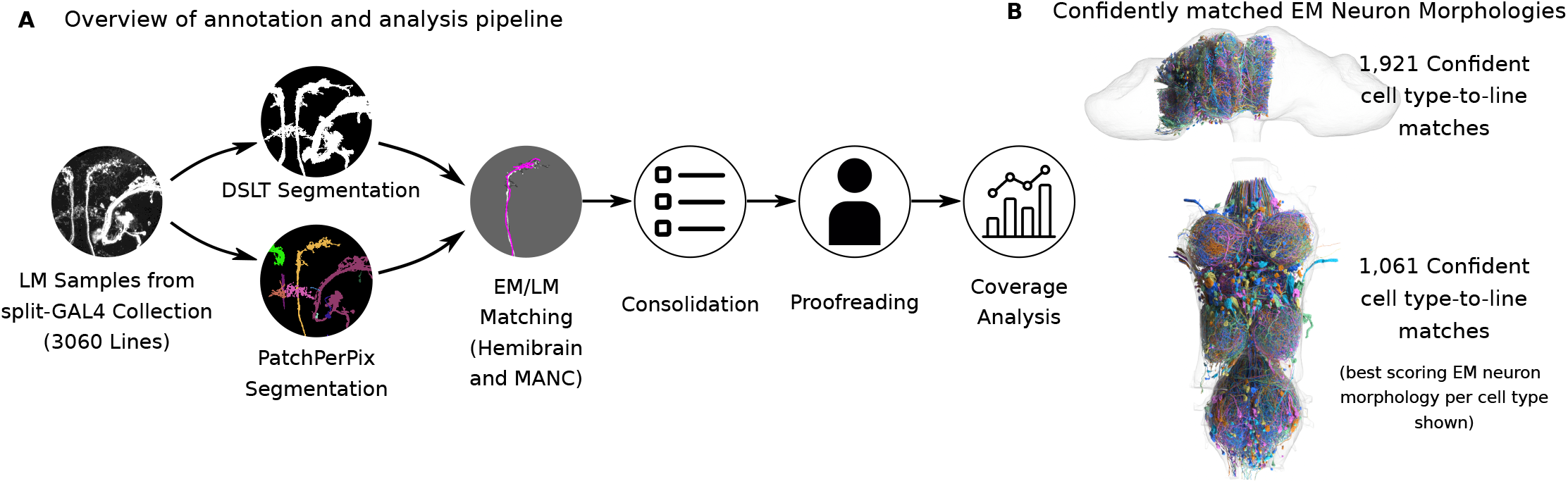
Matching EM-derived cell types to LM samples of the split-GAL4 collection. **A** Schematic of the analysis pipeline. Instance segmentation masks are generated for split-GAL4 samples using PatchPerPix (***Mais et al., 2020***) and DSLT (***Kawase et al., 2015***), and used to identify corresponding neuronal morphologies in EM datasets (hemibrain, ***Scheffer et al., 2020***; MANC, ***Takemura et al., 2024***) via PatchPerPixMatch (***Mais et al., 2021***). Results across same-line samples and segmentation methods are consolidated, and summary visualizations are generated for manual assessment of each split-GAL4 line. A subsequent coverage analysis estimates how many EM-defined cell types are represented in the split-GAL4 collection. **B** Overview of matched EM cell types. We manually proofread ∼370,000 high-scoring automated matches, yielding ∼3,000 high-confidence cell type-to-line matches. For each cell type-to-line match, we show the top-ranked EM neuron morphology, illustrating broad coverage across the searched hemibrain and VNC regions.

## Results

### Updates to annotated cell types

We systematically identified known neuronal morphologies from EM datasets (hemibrain, ***Scheffer et al., 2020***; MANC, ***Takemura et al., 2024***) within LM acquisitions of the split-GAL4 collection using a semi-automated EM/LM matching and proofreading pipeline (Figure 1**A**). We focused on the repeated (20x) imaging used to validate the cell-type-specific lines in ***Meissner et al. (2025***). We proofread 1,614 lines for the brain and 1,310 lines for the VNC, sampled randomly from the collection. In total, we proofread 1,772 unique lines, of which 1,152 were annotated for both the brain and the VNC, 462 only for the brain, and 158 only for the VNC. The brain and VNC were proofread and analyzed independently after line sampling, resulting in asymmetries in line selection between the two (see Methods: Manual Annotations). A small number of additional manually-selected lines were annotated for the purposes of pipeline development and blinded proofreading evaluation. In total we evaluated ∼370,000 automated matching results to generate the reported annotations. In addition, we incorporated annotations from recent studies (***Stürner et al., 2025; Cheong et al., 2026; Ehrhardt et al., 2023; Shuai et al., 2024; Zung et al., 2025***). We provide the full set of available annotations in an updated version of the supplement from ***Meissner et al. (2025***) (see Supplemental File [1]). The NeuronBridge neuron search website (https://neuronbridge.janelia.org ; ***Clements et al., 2024***) has also been updated to display EM-matched cell type information where available.

For proofreading, automated search results were scored by a team of annotators experienced in assessing EM and LM neuron morphologies. Matches were classified as *Confident, Candidate*, or *False*, with the constraint that each cell type observed in LM could be associated with at most one *Confident* EM match. A *Confident* label indicates that the annotator judged the EM/LM match to be a clear positive correspondence, whereas *Candidate* refers to uncertain cases that could represent true matches but lacked sufficient or unambiguous morphological evidence. In some cases, multiple similar cell types (or subtypes, where defined) were all scored as *Candidate* matches and subsequently grouped into a single *Confident* match. These grouped cases are listed in Supplemental File [4] but were not included in the coverage analysis (see below).

We identified 6,368 Confident and Candidate cell type-to-line matches across the hemibrain and MANC datasets, including 29 cell type-to-line matches that were independently confirmed in both the brain and ventral nerve cord (Figure 2**A**). Restricting the analysis to the 2,953 Confident matches, we assigned cell type annotations to 1,361 lines, covering 1,617 unique cell types. Previously published annotations, collated by ***Meissner et al. (2025***), provide annotation information for 2,667 lines. Because these annotations were compiled from multiple independent studies, they do not follow a single annotation scheme. To enable comparison, we automatically mapped the published annotations to standardized hemibrain and MANC cell types where possible. This mapping was successful for 1,254 of the 2,667 previously annotated lines, yielding 1,606 cell type-to-line matches corresponding to 950 unique cell types. Of the 1,361 lines we annotated, 725 are not among the 1,254 mapped published lines and thus newly receive Confident cell type annotations through our study. The complement 636 of the 1,361 lines we annotated are also represented among the 1,254 mapped published lines. Within this shared set, our annotations confirmed 582 previously reported cell type-to-line matches, whereas 97 are conflicting with our annotations and therefore require further investigation. Moreover, we add 710 novel cell type-to-line matches to the shared set of lines. These comparisons should be interpreted with caution, as differences in naming conventions, annotation granularity, and curation strategies may contribute to the observed discrepancies. A detailed comparison of the annotated cell types is provided in Supplemental File [7].

**Figure 2.**
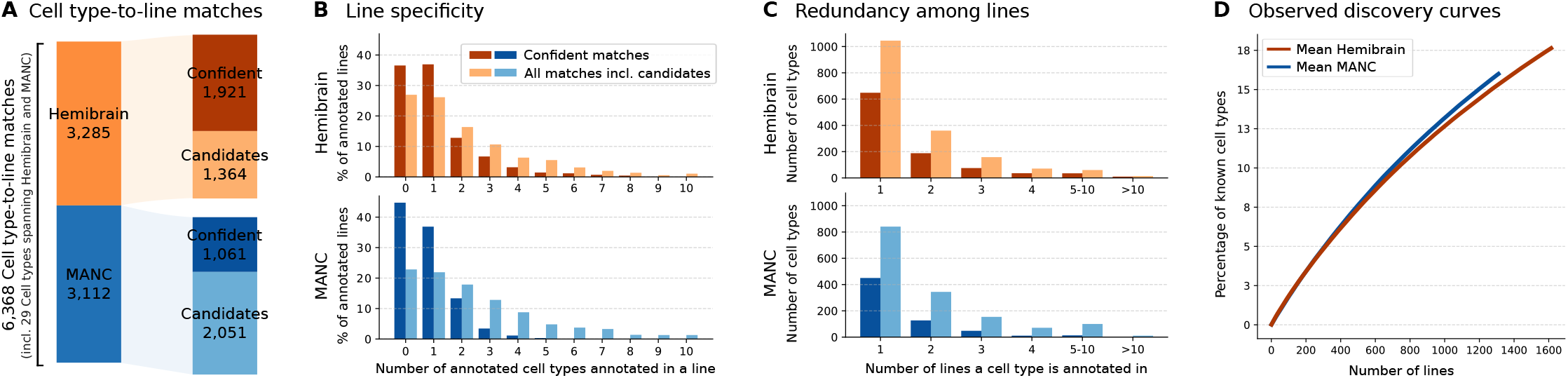
Statistics of cell type annotation updates. **A** Distribution of cell type-to-line matches across the hemibrain and MANC datasets, separated into Confident and Candidate matches. **B** Specificity of the lines: Percentage of annotated lines (y-axis) receiving a certain number of cell type-to-line associations (x-axis), where 100% corresponds to the 1,614 proofread lines for the brain and the 1,310 proofread lines for the VNC. Some annotated lines received no cell types (x=0) because their expression falls outside the examined area—either being specific to the respective other CNS area, or, in the case of the brain, being expressed outside the hemibrain. Other lines lack reliable automated matching results, or only have EM/LM correspondences without an assigned cell type yet. The percentage of lines with no or only a single cell type association decreases when considering *all cell type-to-line matches incl. Candidates*, as Candidate assignments are included even when no Confident cell type-to-line match can be established. **C** Redundancy across lines: Number of cell types (y-axis) that were each associated with a certain number of lines (x-axis). For example, about 200 hemibrain cell types were confidently identified in two lines each. **D** Mean discovery curves for hemibrain and MANC, using Monte Carlo sampling across all Confident annotations. For reference, 5,620 cell types are known in the hemibrain and 4,076 in the MANC.

We calculated the distributions of annotated cell types per line (how specific each line is) and lines per cell type (how much redundancy is present in the split-GAL4 collection) (Figure 2**B/C**). Due to expression outside the examined area or technical reasons (see Methods: PPPM Evaluation and Methods: Manual Annotations), some lines had no matches. On average, each line yielded 1.19 Confident cell types in the brain and 0.8 in the VNC. Including Candidate annotations increases these averages to 2.04 and 2.38 cell types per line, respectively. On average, each Confident cell type was identified in 1.94 lines for the brain and 1.63 lines for the VNC, and including Candidates each cell type was identified in 1.92 and 2.05 for the brain and the VNC respectively. Figure 3 illustrates example EM/LM matches of cell types occurring in multiple lines. Figure 2**D** shows the discovery curve, i.e., the percentage of distinct cell types identified as a function of the number of annotated lines. The curve indicates that the number of newly identified unique cell types continued to increase throughout the annotation effort. See below for further analysis of cell type capture in the collection. Together, these analyses highlight the balance between line specificity and collection coverage. While highly specific lines label few cell types, broad coverage of the nervous system requires many distinct lines, as reflected by the continuously increasing discovery curve.

**Figure 3.**
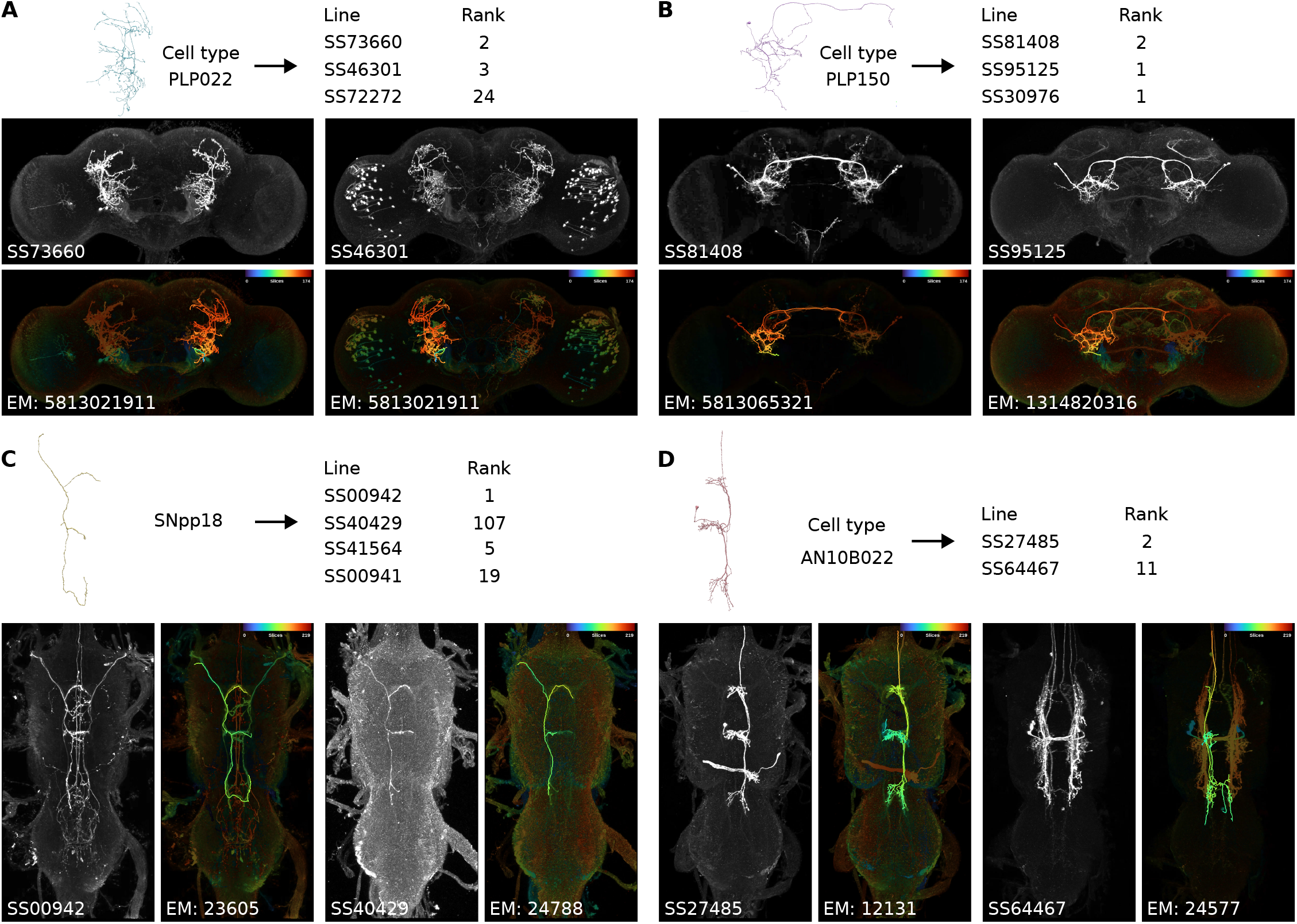
Example cell types identified across multiple lines in the brain (**A**,**B**) and VNC (**C**,**D**). For each LM/EM match, the EM body ID (neuPrint, ***Clements et al., 2020; Scheffer et al., 2020; Takemura et al., 2024; Marin et al., 2024***) is shown on top, alongside a grayscale maximum intensity projection (MIP) of the full expression pattern and the corresponding color depth–encoded MIP with the EM body ID overlaid. The reported rank reflects the automated PPPM cell type ranking, with lower values indicating better matches. In **A**, we confirm the previously published annotation of PLP022 in line SS73660 and identify the same cell type in two additional lines. In **B** and **C**, we newly identify PLP150 and SNpp18 in three and four lines, respectively. **D** shows the newly identified cell type AN10B022 in two lines; in SS64467, it represents a secondary cell type.

We further evaluated the accuracy of our annotation protocol using previously published annotations. In a blinded proofreading assessment, our annotators identified these published matches with miss rates of 12% for the brain and 16% for the VNC (see Methods: Blinded Proofreading Testing). PPPM, whose automated matching results form the basis for proofreading, achieved a median cell type rank of 4 for the hemibrain and 3 for the MANC, indicating that half of the published matches were ranked within the top four or three positions, respectively (see Methods: PPPM Evaluation). Note that our blinded evaluation measures the recovery of previously published matches, which often focus only on the primary cell type. We did not perform an independent inter-annotator agreement assessment of the additional matches identified by our annotators. While we do not expect our judgments to be the final word on specific cell type matches, we view these annotations as a valuable extension to the split-GAL4 collection resource.

### Cell type-based coverage analysis

***Meissner et al. (2025***) estimated the spatial distribution of the split-GAL4 collection based on registered voxels illuminated in images of split-GAL4 lines, introducing uncertainty from details of imaging and foreground/background segmentation. We revisited this question from a different direction, asking how many unique cell types are labeled by individual lines and modeling the contents of the collection as a whole. Our analysis, being based on our annotation efforts as described in the previous section, is restricted to neurons likely to be identifiable in the hemibrain and MANC volumes (***Scheffer et al., 2020; Takemura et al., 2024; Marin et al., 2024***).

We modeled the process of discovering cell types in the split-GAL4 collection using a Monte Carlo simulation, based on empirical observations from a subset of annotated lines. The goal was to estimate the cumulative number of distinct cell types discovered as a function of the number of lines, and to extrapolate this curve to the full collection. Robustness was ensured through Monte Carlo cross-validation and repeated simulations, with final discovery curves reported as the mean and percentile ranges across all models, and analogous analyses performed for individual anatomical regions.

The extrapolated discovery curves are shown in Figure 4**A** and listed in detail in Supplemental Files [2] and [3]. Based on all Confident cell type-to-line matches, we estimate full collection coverage of about 26% of the known cell types in the hemibrain and 28% in the MANC. This corresponds to an estimated mean of 1,445 unique cell types for the brain, with a standard deviation of ±49 cell types and a 95% simulation interval spanning 200 cell types. For the VNC, the corresponding estimate is 1,145±47 unique cell types, with a 95% simulation interval of 182 cell types. If we also include all Candidates on top of Confident matches, coverage increases to about 43% and 57% for the hemibrain and MANC respectively. For the individual regions of the hemibrain, we estimate coverage varying from 23% in the Great Commissure or Lateral Horn to 59% for the Central Complex. The regions with greatest coverage likely result from focused efforts to catalog neurons, suggesting similar efforts could raise coverage elsewhere (***Wolff and Rubin, 2018; Wolff et al., 2025; Nern et al., 2015; Wu et al., 2016; Nern et al., 2025***). In the MANC, the spatial distribution is more uniform, with coverage ranging from 25% to 38% (see Figure 4**B**).

**Figure 4.**
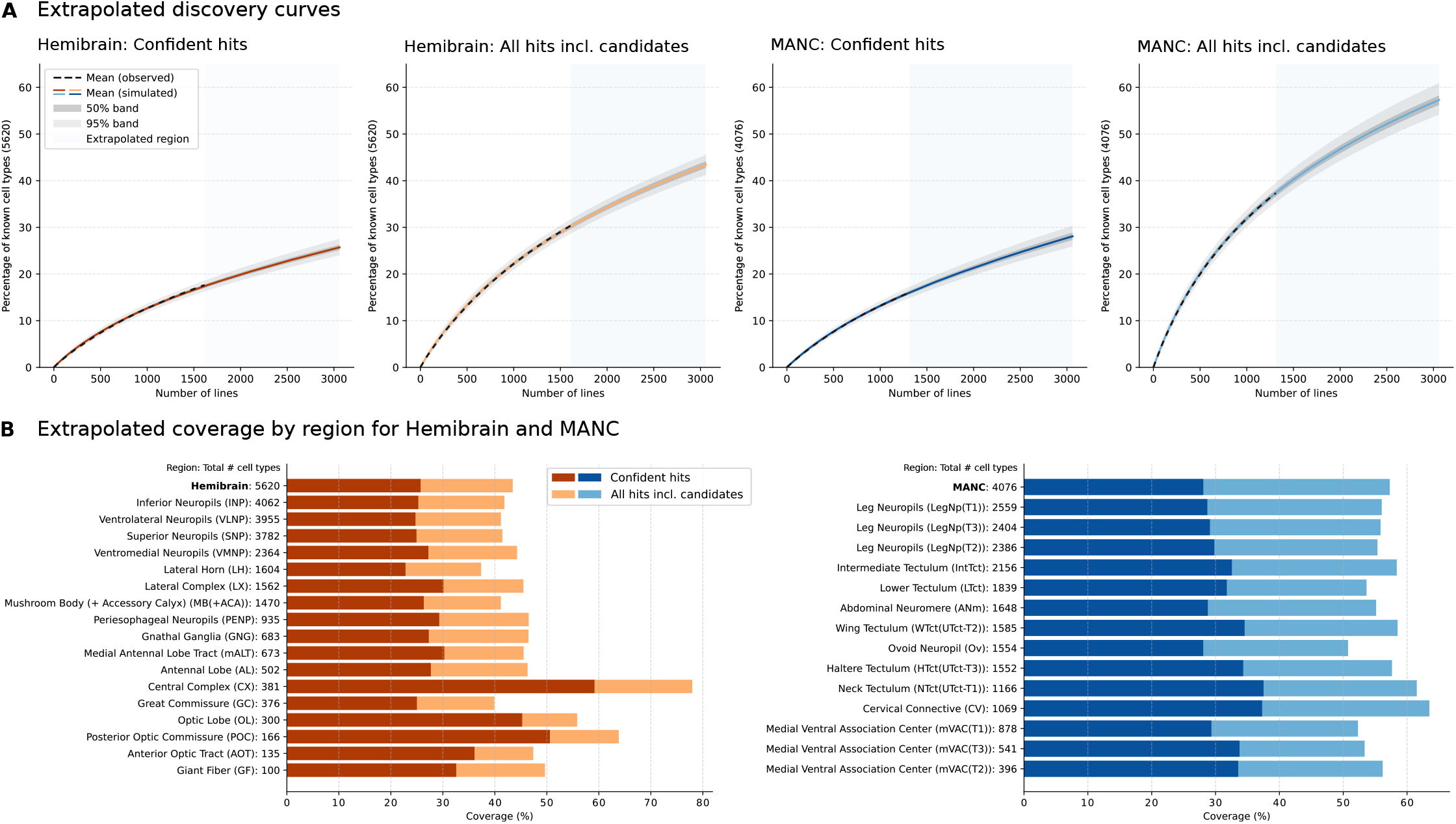
Extrapolated coverage of cell types in the split-GAL4 collection. **A** Discovery curves showing the cumulative fraction of uniquely identified cell types as a function of the number of split-GAL4 lines, for hemibrain and MANC datasets. Results are shown for high-confidence matches and all matches including candidates. The colored solid lines indicate the mean of the respective extrapolated (simulated) discovery curve; shaded bands around the curve show 50% (dark shade, best viewed at >500% zoom) and 95% (light shade) intervals of simulation runs; the dashed lines show the respective observed discovery curve; and the gray region of the plot denotes extrapolation beyond the observed data. We estimate that the split-GAL4 collection covers about 26% of the known hemibrain cell types and 28% of the known MANC cell types based on Confident cell type-to-line matches. Including Candidate matches, these estimates increase to 43% and 57%, respectively. **B** Estimated cell type coverage by anatomical region for hemibrain (left) and MANC (right). Regions follow definitions at https://neuprint.janelia.org, and some hemibrain volume regions (such as Optic Lobe) contain only a portion of the full brain regions. Bars indicate the fraction of cell types covered by Confident matches (dark shade) and all matches including candidates (light shade).

For each model, we evaluated simulation performance using the distance between the mean predicted number of unique cell types and the observed number of unique cell types across all annotated lines. Assuming that this prediction error corresponds to a fixed multiple of the standard deviation of the simulated values, we extrapolated it to the full set of lines in the split-GAL4 collection (see Methods: Validation). Based on this approach, the estimated prediction error for the brain is 23±16 for Confident cell type-to-line matches and 23±19 when also including Candidates. For the VNC, the corresponding predicted errors are 15±12 for Confident cell type-to-line matches and 24±20 when also including candidates. Note that the estimated prediction errors and the reported percentile ranges (shown in Figure 4) quantify different sources of uncertainty: Prediction errors are estimated separately for each model based on left-out annotated lines, whereas the reported percentile ranges are calculated across simulations from all models. Accordingly, the latter are expected to be larger because they additionally reflect variability between models.

We expect our estimates based on the Confident cell type-to-line matches to somewhat under-estimate the real coverage, as retrieval of all expressed cell types per line was not always possible. Thus, the coverage analysis presented here reflects discovery achievable *through our pipeline*, not necessarily true biological coverage, and is subject to uncertainties such as matches missed during automated matching or manual proofreading. Whether estimates based on all cell type-to-line matches, including Candidates, more accurately reflect the true coverage remains uncertain. Nonetheless, the estimated CNS coverage of the split-GAL4 collection is substantially lower when assessed at the level of cell types compared to evaluations based on labeled voxels in ***Meissner et al. (2025***).

## Discussion

We extend the split-GAL4 driver collection presented by ***Meissner et al. (2025***) with additional EM-derived cell type annotations and make these annotations readily accessible through the Neuron-Bridge search tool (https://neuronbridge.janelia.org). Leveraging these annotations, we provide a complementary estimate of the collection’s CNS coverage based on identified neuronal cell types. Together, these contributions increase the utility of the split-GAL4 collection for identifying driver lines, interpreting experimental results, and guiding future studies of Drosophila neural circuits.

Our estimate of CNS coverage based on confidently matched cell types is substantially lower than the previously reported voxel-based estimate. However, these values are not directly comparable, as they quantify two related but distinct aspects of the split-GAL4 collection. The voxel-based analysis of ***Meissner et al. (2025***) measures the fraction of voxels labeled by one or more split-GAL4 lines, whereas our analysis estimates the fraction of distinct neuronal cell types represented in the collection. A small number of broadly arborizing neurons can occupy a large fraction of anatomical space, while many spatially restricted cell types contribute relatively little. In addition, the limited resolution of light microscopy introduces partial volume effects, whereby imaged voxels are larger than the underlying neuronal processes, inflating their apparent occupied volume, while conversely multiple neurons may contribute to the same image voxel. The voxel-based regional analysis also includes the passage of neuronal processes through an anatomical compartment, rather than being based on synaptic innervation of the compartment.

Our approach relies on a more complex analysis pipeline, and the estimates depend on the quality of automated EM/LM matching, manual proofreading, and the statistical model used for extrapolation. Although coverage could, in principle, be estimated at the level of individual EM body IDs, we chose to use EM-defined cell types, as they provide a useful means to subdivide the CNS into related units. However, cell type definitions continue to evolve and can vary in terms of size, nomenclature, internal divisions, and instantiations across individual organisms, making our mapping provisional. Furthermore, driver expression within a cell type often does not mean fully labeling all cells making up the type, so the count is likely to trend high compared to measuring individual neurons. Despite these challenges, we believe that our cell type analysis provides a more biologically meaningful and easily interpretable approximation of the fraction of neuronal populations that can be specifically targeted by the current split-GAL4 collection.

Beyond directly facilitating physiological and behavioral studies as well as enabling the coverage analysis presented here, the curated annotations also provide a valuable resource for future improvements in automated EM/LM matching. PatchPerPixMatch effectively prioritizes candidate EM/LM matches for manual review, but as with other approaches has a false positive rate that precludes fully automated one-to-one matching (***Mais et al., 2021***). We expect future EM/LM matching methods to increasingly replace hand-engineered components with learned representations, for example using graph neural networks or related deep learning approaches. The ∼370,000 manually annotated search results represent a highly valuable resource for this purpose as they include not only correct matches but also high-ranking false positives from challenging cases – data that was previously unavailable. While these annotations are not yet organized as a standardized benchmark with predefined training and evaluation splits, establishing such a benchmark together with baseline methods would enable rigorous comparison of automated matching algorithms and represents an important direction for future work. As EM connectomes continue to provide increasingly complete cell type catalogs, robust methods for linking these cell types to genetic driver collections will become ever more important. By expanding the available cell type information for the split-GAL4 collection and providing a large set of manually reviewed annotations, this work lays a foundation for future efforts to bridge connectomics and genetic tools.

## Methods and Materials

### EM/LM Neuron Matching and Visualization for Efficient Annotation

We describe the EM/LM matching procedure used to identify known neuronal morphologies from the hemibrain (***Scheffer et al., 2020***) and MANC (***Takemura et al., 2024***) EM datasets within the LM acquisitions of the ***Meissner et al. (2025***) split-GAL4 line collection. We followed a two-stage pipeline in which instance segmentations of the LM acquisitions are first obtained using two complementary methods and subsequently used for EM/LM matching with *PatchPerPixMatch* (***Mais et al., 2021***). We randomly sampled split-GAL4 lines from the collection that exhibited sparse expression patterns (corresponding to Line Quality 1 and 2 in ***Meissner et al. (2025***); 2,999 of 3,060 lines). For these lines, we performed *PatchPerPixMatch* on both the brain and VNC LM acquisitions. The resulting automated EM/LM matches were then distributed among annotators for manual evaluation, with the goal of annotating as many lines as possible. Below, we detail segmentation and matching methods, as well as how we compile and visualize the resulting matches to facilitate manual annotation.

#### LM Neuron Segmentation

To improve downstream matching quality, we applied two complementary light microscopy (LM) neuron segmentation methods: *PatchPerPix* (***Mais et al., 2020***) and *Direction-Selective Local Thresholding* (DSLT; ***Kawase et al., 2015***).

First, we trained the standard *PatchPerPix* architecture, referred to as ppp in ***Mais et al. (2020***) on the FISBe dataset (***Mais et al., 2024***). The three-channel FISBe data were converted to grayscale image volumes, and training was performed on both the original resolution (0.44 *µ*m^3^ per voxel) and a 2× downsampled version. All input data were normalized to zero mean and unit variance for both training and inference. The model was optimized using a cross-entropy loss with a batch size of 2. We employed a 3-level 3D U-Net (***Ronneberger et al., 2015***) with 2× downsampling and upsampling operations. The network was initialized with 20 feature maps, tripled at each downsampling level. For each voxel, the model predicts patches of size 7 × 7 × 7 from which we puzzle together the instance masks. Second, we used DSLT-based segmentation masks provided by ***Meissner et al. (2025***). Since DSLT produces foreground rather than instance segmentations, connected components were used as proxies for individual neuron instances.

Because both methods operate on grayscale images of full expression patterns, adjacent neuronal processes may merge into a single segment. The two approaches exhibit complementary failure modes: PatchPerPix tends to falsely split processes, whereas DSLT tends to include surrounding background regions in the segmentation of dense dendritic structures, resulting in the merging of distinct branches and poor-quality skeletons. In the following, we refer to instance masks from both approaches collectively as *(LM) fragments*.

#### EM/LM Neuron Matching and Visualization of Results

Automated EM/LM matching was conducted using *PatchPerPixMatch* (***Mais et al., 2021***), a combinatorial solver that takes a target neuron morphology (from EM) and determines whether it is present among a set of other morphologies (fragments from a LM acquisition) considering potential false merges and splits. We followed the procedure described in the original work, except that we applied it to two different LM segmentation bases and replicated grayscale values across three channels to enable use of the color penalty term without modification. We used EM neuron morphologies from the hemibrain (v1.2.1) and MANC (v1.2.1) datasets.

The matching was carried out for each pair of split-GAL4 sample and EM neuron morphology (referred to as an EM body ID). Because each split-GAL4 line was imaged in at least two samples (female and male, 20x CsChrimson ‘rescreening’ images), we consolidated matching results across samples as well as across both segmentation-based runs (PatchPerPix and DSLT). For each split-GAL4 line, the top matches were then compiled into a ranked list of candidate EM neurons. To facilitate efficient manual annotation, we generated PDF visualizations summarizing the top-scoring matches, organized as follows:

- Previously published cell type or EM body ID annotations are presented at the top, when available, to provide high-confidence reference points and prevent published annotations from being missed due to low automated matching scores.
- Matches were grouped by EM cell type, where available.
- For each cell type, only the top five EM body IDs were included.
- Matches were sorted by spatial proximity using hierarchical clustering (***Pedregosa et al., 2011; Sokal and Michener, 1958***) of EM neuron bounding boxes, with up to 60 matches shown per cluster.

An example PDF visualization is provided in Supplemental File [5].

#### Neuron Exclusion Criteria

To avoid small or ambiguous fragments in the high-scoring matching results, we applied the following empirically determined neuron exclusion criteria. For the hemibrain, we excluded all EM body IDs that are smaller than 200 million voxels in the EM segmentation, have no identified cell type, *and* are cropped at the border of the acquired volume. For the MANC dataset, we applied a broader filter across the entire dataset, excluding EM body IDs that were smaller than 100 million voxels in the EM segmentation and had no identified cell type. The voxel-count thresholds above refer to the neuron *size* property from the neuprint API (***Clements et al., 2020***)—the number of voxels comprising each segmented EM body—reported at the native 8 × 8 × 8 nm isotropic voxel resolution used for both the hemibrain and MANC EM volumes. The mean body size across the complete dataset was 96 million voxels for the hemibrain and 194 million voxels for MANC. Importantly, this means that large neurons are kept, even without an assigned cell type, as they may still yield high-quality matches. We note that these dataset-specific filtering criteria result in subsets of EM neuronal morphologies used for automated matching that differ in their distribution of EM body IDs with and without assigned cell type annotations. In particular, the filtered hemibrain dataset contains more EM body IDs without assigned cell types than the MANC dataset (Figure 5**B**), which is also reflected in the manually annotated matches. Note that this filtering did not exclude any known cell types from either the hemibrain or the MANC dataset — all such cell types remained represented in the filtered set. For LM fragments, we filtered search results at the end of the matching for both hemibrain and MANC when the selected skeletons had less than 200 pixels in total.

**Figure 5.**
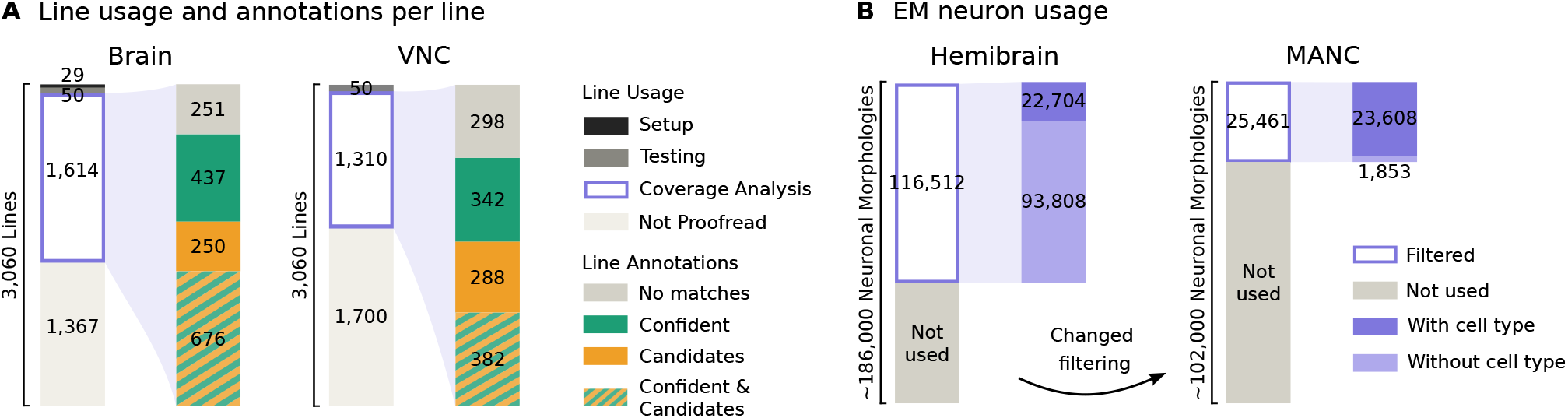
Overview of line and EM neuronal morphology usage. **A** Lines are categorized into four usage groups: pipeline setup, blinded proofreading testing, coverage analysis (randomly sampled), and lines not proofread. Within the lines sampled for coverage analysis, we further indicate the number of lines with Confident and/or Candidate annotations, as well as lines with no identified matches. Note that we include all proofread EM/LM matches here, even those for EM body IDs not yet assigned a cell type. **B** Usage of EM neuronal morphologies from the hemibrain and MANC datasets after dataset-specific filtering, highlighting differences in the inclusion of EM body IDs with and without an assigned cell type.

Identical filtering procedures were applied in both PatchPerPixMatch and the coverage analysis. Coverage analysis was performed exclusively at the *cell-type* level (see Methods: Manual Annotations), but all EM body ID matches without assigned cell types are provided in Supplemental Files [2] and [3] as a resource.

#### PPPM Evaluation

To assess the matching accuracy of PPPM, we used Confident matches, and evaluated the rank at which they were retrieved by PPPM. Because our primary focus is on cell types—and each cell type may correspond to multiple EM body IDs—we performed the evaluation at the cell-type level. Ranks were determined based on the best matching score between an EM body ID and the query sample. Here, top-ranked matches correspond to the highest matching scores and appear at the top of the ranked list, whereas lower-ranked matches have lower matching scores and appear further down the list.

Table 1 summarizes the distribution of PPPM cell-type ranks for previously published annotations that we confirm and for all Confident cell type-to-line matches identified in this study. Note that the number of previously published annotations confirmed by us differs between this evaluation and the Results (Section Updates to annotated cell types) for two reasons: (1) this evaluation uses the annotations as published in ***Meissner et al. (2025***), whereas the Results incorporate updates from other recent studies (***Stürner et al., 2025; Cheong et al., 2026; Ehrhardt et al., 2023; Shuai et al., 2024; Zung et al., 2025***); and (2) annotators were provided with all annotations, not just those matched automatically, as higher-level annotations, abbreviations, and alternative terminology may still be informative. For the previously published matches, PPPM achieved a median cell-type rank of 4 for the hemibrain and 3 for the MANC, indicating that 50% of published matches were retrieved within the top four and top three positions, respectively. At the 75th percentile, 75% of published matches were retrieved within the top 12 positions for the hemibrain and the top 9 positions for the MANC. Compared with the previously published matches, the Confident cell type-to-line matches from this study were often retrieved at lower positions in the ranked list, indicating lower matching scores. This likely reflects the inclusion of secondary or tertiary cell types, as well as more challenging cases for PPPM. PPPM outputs a maximum of 170 EM body IDs per query for manual review. The exact composition of this output depends on the number of EM body IDs assigned to each cell type and the number of matches within each regional cluster. Consequently, true matches ranked beyond this cutoff are not included in the PDF visualization and may be missed during manual review. Previously published annotations are explicitly placed at the top of the PDF and are therefore retained regardless of their rank. Without this prioritization, PPPM would have missed approximately 5% of published annotations for the MANC dataset and fewer than 5% for the hemibrain.

**Table 1.** Summary statistics of PPPM cell-type ranks for the hemibrain and MANC datasets, computed separately for two groups: previously published annotations confirmed by our analysis, and the full set of manually annotated *Confident* cell type-to-line matches from this study (which subsumes the first group). Ranks are defined by the highest-scoring EM body ID within each cell type, with lower ranks indicating better matching. Most matches are recovered among the top-ranked cell-type candidates (median rank 3 to 5; 75th percentile rank 9 to 16), while a small fraction are retrieved much further down the list (95th percentile rank up to 132). Please see Methods: PPPM Evaluation for more interpretation.

|  | Hemibrain |  | MANC |  |
| --- | --- | --- | --- | --- |
|  | Published | All Confident | Published | All Confident |
| Num matches | 560 | 1,921 | 142 | 1,061 |
| Median rank | 4 | 5 | 3 | 4 |
| 25th percentile rank | 1 | 2 | 1 | 1 |
| 75th percentile rank | 12 | 16 | 9 | 14 |
| 95th percentile rank | 69 | 64 | 132 | 49 |
| Min rank | 1 | 1 | 1 | 1 |
| Max rank | 327 | 327 | 1,035 | 1,035 |
| Mean rank | 14.98 | 15.26 | 29.5 | 13.7 |

Please note that we do not evaluate false positives here because the method returns a ranked list of candidates for manual proofreading rather than making final predictions. Since the number of true matching cell types differs between lines and is generally not exactly known (not annotated), no objective cutoff exists for converting the ranking into positive and negative predictions. Any false-positive estimate would therefore depend on an arbitrary threshold rather than on the underlying matching performance.

Beyond the quantitative evaluation of published and newly identified EM/LM matches in the provided ranked lists above, we identified typical failure cases in lines for which PPPM did not yield any convincing (Confident or Candidate) matches (Figure 5**A**). No matches were identified for 251 lines in the brain and 298 lines in the VNC. Besides brain- or VNC-specific expression patterns, the automated matching procedure may fail in case of poor segmentation of dim neurons or neuron bundles, and expressed neurons lying outside the hemibrain volume (e.g., optic lobes or ascending neurons); examples are shown in Figure 6.

**Figure 6.**
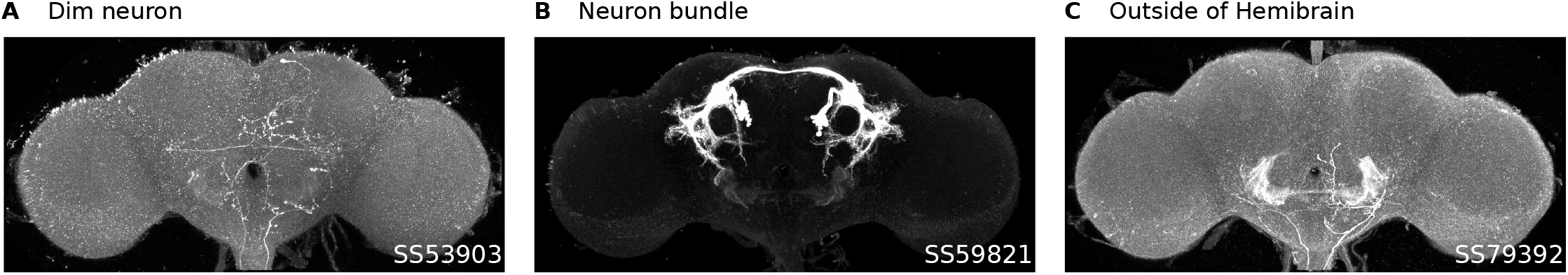
Representative failure cases. **A** Dimly labeled neurons often produce incomplete or fragmented segmentation masks. **B** Neuronal bundles frequently yield merged, dilated masks that cannot be reliably skeletonized for morphology matching. **C** Neurons located outside the hemibrain volume may only be represented by small cropped EM morphologies, which can also prevent successful matching.

### Manual Annotations

To identify true EM/LM correspondences among the automated matching results, we manually evaluated the top-scoring candidates per line (see Methods: EM/LM Neuron Matching and Visualization of Results). Five trained annotators, all experienced in interpreting EM and LM neuronal morphologies, labeled each PPPM match shown in the PDF visualizations as *Confident, Candidate*, or *False*. A *Confident* label indicates that the annotator judged the EM/LM match to be a clear positive correspondence. *Candidate* refers to uncertain cases that could represent true matches but lacked sufficient or unambiguous morphological evidence, while *False* indicates a confident negative judgment.

Annotation criteria included the relative positions of neuronal processes and cell bodies, the overall neuronal morphology, and whether morphological similarities were consistent along the full extent of the structure. During annotation, each cell type observed in LM was constrained to have at most one *Confident* EM match. If multiple similar cell types (or subtypes) were plausible but none met the Confident threshold, all corresponding EM candidates were labeled as *Candidate*. In general, each line was assessed by a single annotator; however, in cases of uncertainty, a second annotator reviewed and verified the initial judgment. To ensure procedural consistency across annotators, annotation guidelines were documented (provided in Supplemental File [5]), and all annotators underwent an initial training session. In post-processing, similar subtypes were grouped into a single *Confident* cell type when the collective evidence supported a unique identity; this grouping was not used in the coverage analysis but is provided as additional annotation (see Supplemental File [4]).

In Figure 5**A**, we summarize the number of lines that were manually proofread for the brain and the VNC, and whether Confident and/or Candidate matches were identified (considering all matches, i.e., also including EM/LM matches for EM body IDs without an assigned cell type). Out of 3,060 split-GAL4 lines, hemibrain matches were scored for 1,693 lines and MANC matches for 1,360 lines. Note that the sets of lines annotated for the brain and VNC are not identical, although they partially overlap. This is because proofreading and analysis of the brain and VNC were performed independently and within a fixed time budget, with the goal of annotating as many lines as possible. As a result, the annotation is biased toward sparser lines, as some of the more difficult, denser lines were left incompletely proofread within the available time. In total, we spent approximately 85 annotation days on the lines used for the coverage analysis, requiring on average 16 minutes per PDF (brain or VNC). In Figure 3, we show example cell types identified across multiple lines, and in Figure 7 we show example lines with multiple identified cell types. Note that the automated search results were manually proofread for each EM body ID-to-line match. For the coverage analysis, the proofread matches were consolidated at the cell type level, which are provided in Supplemental Files [2] and [3]. All individual EM body ID-level labels, including automated matches labeled *False*, are available in the coverage analysis repository (see Data availability).

**Figure 7.**
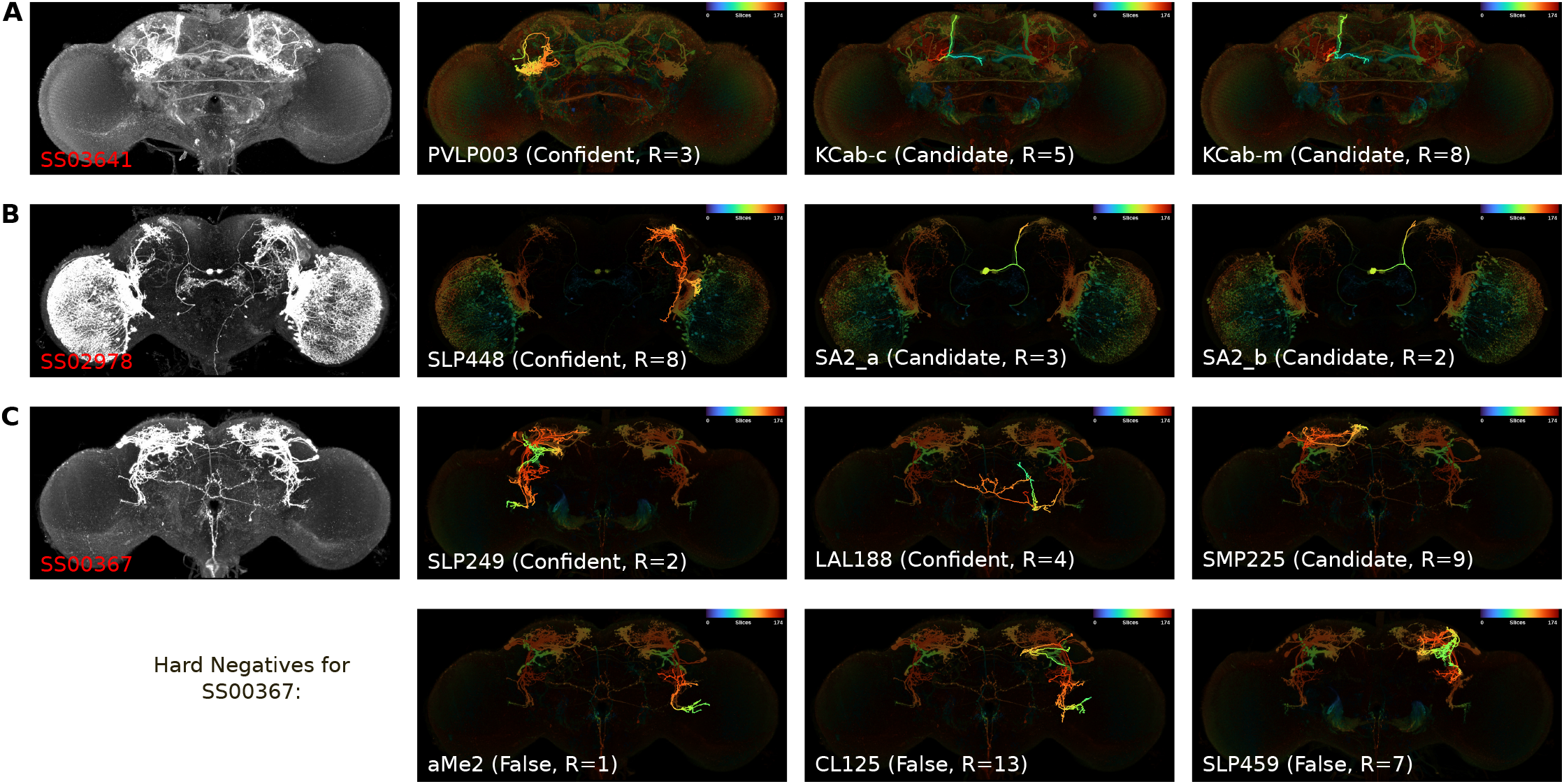
Example lines with multiple cell types identified. First column shows full expression pattern of given line, and other columns show color depth MIP of expression pattern with corresponding EM neuron overlaid. Line name is given in red, cell type information in white with corresponding manual annotation (Confident, Candidate, False) and automated PPPM cell type rank (R). **A** shows line SS03641 with cell type PVLP003, previously published and confirmed by our annotation, and KCab as a secondary cell type newly identified here. While the subtypes KCab-c, KCab-m, and KCab-s (not shown here) were annotated as Candidate cell type-to-line matches, we are confident that cell type KCab represents a true match for line SS03641 (see sub cell type grouping in Supplemental File [4]). **B** For line SS02978, we newly identified cell types SLP448 and SA2. While SA2_a and SA2_b were initially annotated as Candidate matches, we classify SA2 as a Confident match. **C** For line SS00367, we confirmed the previously published match to SLP249 and newly identified LAL188 as a Confident match. SMP225 (together with SMP230 for the same cell, not shown) was annotated as a Candidate match. The second row shows high-ranking automated search results that were classified as False by our annotators.

#### Blinded Proofreading Testing

To validate our five-person manual proofreading process, we performed a single-blinded evaluation of our annotations (i.e., the classification of automated matching results as *Confident, Candidate*, or *False*) against previously published EM/LM annotations collected in ***Meissner et al. (2025***). For this purpose, we generated PDF visualizations in which previously published annotations were presented according to their ranking by the automated matching algorithm rather than being placed at the top of the list. We verified that all previously published annotations were included. Each annotator evaluated 10 lines for hemibrain matches and 10 lines for MANC matches, and our annotations were compared to the published annotations (Supplemental File [6]). As several lines had multiple published annotations, the total number of published annotations was 56 for the hemibrain and 82 for the MANC. Upon closer inspection, our most experienced annotator (C.M.) determined that 6 of the 56 (11%) hemibrain annotations and 13 of the 82 (16%) MANC annotations were likely incorrect; these were excluded from the quantitative evaluation. Of the remaining 50 hemibrain and 69 MANC annotations, our team correctly identified 44 hemibrain and 58 MANC matches, corresponding to miss rates of 12% and 16%, respectively.

### Coverage Analysis

This section describes the probabilistic model used to estimate the total number of unique cell types labeled by the full collection of split-GAL4 lines described in ***Meissner et al. (2025***), based on annotations from roughly half of the available lines. This problem is related to species richness estimation (***Schmitz and Rahmann, 2025***), where the total number of distinct classes is inferred from a subsample. Classical estimators such as Chao1 (***Chao, 1984, 1987***) and jackknife (***Burnham and Overton, 1979***) are known to underestimate richness in highly heterogeneous populations with many unobserved classes. More complex parametric or upsampling-based methods require assumptions about the underlying distribution or sampling process, like independent and identically distributed sampling of individual observations, that are not satisfied in our setting. In our case, the observations correspond to cell types identified across split-GAL4 lines, where each line can be viewed as a “package” containing a subset of cell types, similar to the coupon collector problem (***Stadje, 1990***). However, the number of cell types observed per line varies, and each cell type can be observed at most once within a given line, corresponding to sampling without replacement within each package. While a direct, closed-form, probabilistic formulation of the discovery process is formally possible, it becomes computationally intractable due to dependencies between observations, as evaluating the full likelihood would require enumerating all possible sampling sequences. We therefore model the process of cell type discovery using Monte Carlo simulation, which approximates the underlying distribution without requiring explicit enumeration of dependent sampling outcomes. In the following, we define the cell type discovery curve and describe its empirical and simulated formulations. We then explain how empirical distributions are derived from the annotated data and used to parameterize a simulation model that extrapolates the expected number of discovered cell types across the full split-GAL4 collection. The analysis was performed separately for the brain and VNC.

#### Definition of Cell Type Discovery

We quantified cell type discovery by modeling the cumulative number of unique cell types identified as additional split-GAL4 lines were sampled, yielding the *cell type discovery curve*. Let *L* denote the set of split-GAL4 lines and *C* the set of known cell types. We define the mapping

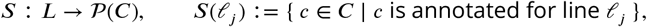

where *P*(*C*) denotes the power set of *C* (i.e., the set of all subsets of *C*), and *S*(*l*_*j*_ ) is the set of cell types expressed in line *l*_*j*_ . Given a line ordering *l*_1_, …, *l*_|*L*|_ of *L*, we define the *discovery function d*(*j*) as the cumulative number of distinct cell types discovered after considering the first *j* sampled lines:

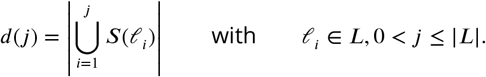

More generally, for any subset *L*^′^ ⊆ *L*, we define 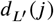 analogous to *d*(*j*), but with *l*_*i*_ ∈ *L*^′^, 0 < *j* ≤ |*L*^′^|, so that *d*(*j*) ≡ *d*_L_ (*j*) is the special case *L*^′^ = *L*. This generalization allows us to apply the discovery function to any subset of lines, such as the subset of annotated lines or lines used for model fitting. In the next two sections, we distinguish two variants of this discovery function: an *empirical* discovery curve 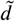, computed directly from the observed annotations *S*(*l*_*j*_ ) for a set of annotated lines, and a *simulated* discovery curve 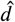, in which we instead simulate lines in terms of their expressed cell types, drawn from a probabilistic model whose parameters are fit to the empirical annotations.

#### Empirical Cell Type Discovery

Let *L*_ann_ ⊆ *L* denote the set of annotated split-GAL4 lines. To obtain a single realization *r* (i.e. a specific fixed ordering of lines) of the empirical discovery curve 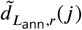, we sequentially sample lines *l*_*j*_ without replacement from the annotated set of lines (*L*_ann_). Because the shape of the discovery curve depends on the ordering of the sampled lines, we repeated this procedure using Monte Carlo sampling with *R*_emp_ = 500 random orderings. The empirical discovery curve was then estimated as the mean

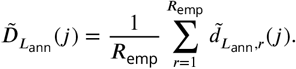

The empirical discovery curves for all Confident cell type-to-line matches for the brain and the VNC are illustrated in Figure 2**D**.

#### Simulated Cell Type Discovery

To extrapolate the cell type discovery curve to the full split-GAL4 collection of |*L*| = 3060 lines, we modeled the discovery process using a Monte Carlo simulation based on empirical observations from the annotated lines. In each realization *r* of the simulation, we sequentially generated, for each position *j* ∈ {1, …, |*L*|}, a simulated set of expressed cell types *Ŝ*_*j*_ ∈ *P*(*C*). Unlike *S*(*l*_*j*_), which denotes the observed annotation for an actual line *l*_*j*_ ∈ *L, Ŝ*_*j*_ is a random variable drawn from a probabilistic model and does not correspond to any specific line in *L*. The simulation of *Ŝ*_*j*_ consisted of two stochastic components:

##### (1) Sampling the Number of Expressed Cell Types

We sampled the number of cell types expressed at position *j*, denoted *k*_*j*_ ∈ ℕ, from the empirical distribution

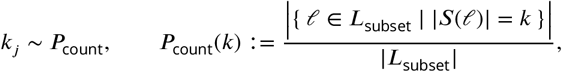

where *P*_count_ is derived from a subset of the annotated lines, denoted *L*_subset_ and defined in Model Fitting. Figure 2**B** shows the distribution of the number of expressed cell types for all annotated lines.

##### (2) Sampling Cell Type Identities

To simulate which specific cell types are expressed at position *j*, we defined a Laplace-smoothed empirical distribution over known cell types:

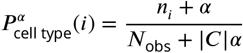

where:

- *n*_*i*_ := |{ *l* ∈ *L*_subset_ ∣ *c*_*i*_ ∈ *S*(*l*) }| is the number of lines in *L*_subset_ that annotate cell type *c*_*i*_,
- 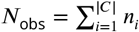 is the total number of observed cell type occurrences,
- |*C*| is the number of known cell types,
- *α* > 0 is the Laplace smoothing parameter, which controls the probability mass assigned to unobserved cell types.

As with *P*_count_, the frequencies *n*_*i*_ and *N*_obs_ are derived from a subset of the annotated lines *L*_subset_ (see Model Fitting for subset definition). We then sampled *Ŝ*_*j*_ without replacement:

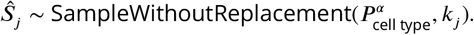

Note that we sampled without replacement, as a single line cannot express the same cell type more than once. To capture sampling variability, we compute multiple realizations *r* = 1, …, ***R*** of the simulated discovery curve over the full line collection, denoted 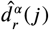, yielding the mean simulated discovery curve

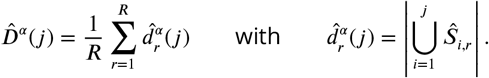

#### Model Fitting

The only tunable parameter in our simulation model is the Laplace smoothing constant *α*, which determines the probability mass assigned to previously unobserved cell types and thereby controls the rate at which new cell types are discovered. To fit and evaluate *α*, we randomly partitioned the annotated lines *L*_*ann*_ into a training set *L*_*train*_ and a held-out set *L*_*held*−*out*_ = *L*_*ann*_ *L*_*train*_, which is not used for fitting. We further sampled a random subset *L*_*ubset*_ ⊂ *L*_*train*_, used exclusively to derive the empirical distributions required by the simulation model as described above. The complete training set *L*_train_, in contrast, served to compute the target empirical discovery curve against which *α* was optimized.

We optimized *α* using Brent’s derivative-free one-dimensional optimization algorithm (***Brent, 1973; Forsythe et al., 1977***), with the objective of identifying the value of *α* that best reproduced the empirical discovery curve 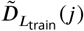 of the training set *L*_train_. At each optimization step for *α*, we averaged multiple realizations of the simulated discovery curve 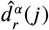 to obtain 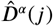 as defined above with *R* = *R*_opt_ = 100. This mean simulated curve was compared with the mean empirical discovery curve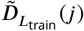, resulting in the squared-error loss

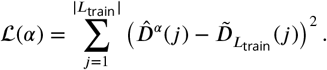

The optimal smoothing parameter was then obtained as

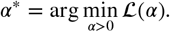

We randomly assigned 75% of annotated lines to *L*_train_, with the remaining 25% held out from fitting (*L*_*held*−*out*_). The held-out lines are used only to test the extrapolation performance of the model (Validation). Two thirds of *L*_train_ were then sampled to form *L*_subset_, used to derive the empirical distributions above. This fitting procedure was performed independently for the brain (1,614 annotated lines) and ventral nerve cord (VNC; 1,310 annotated lines) datasets. Please see Algorithm 1 for an outline of the simulation and Figure 8**A** for an overview of how different sets of lines and their resulting discovery curves are used for model fitting.

**Figure 8.**
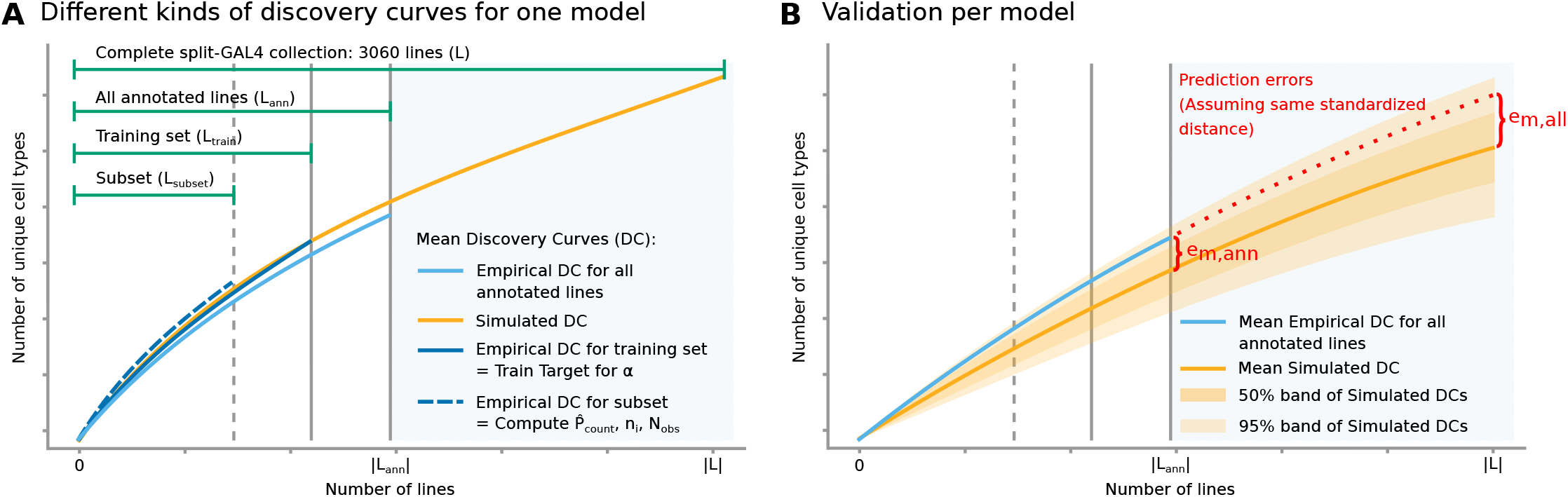
Overview of the training and validation procedure for a **single probabilistic model** *m*. **A** Annotated lines are partitioned into training set and held-out lines, and a subset of the training set is used to derive the empirical distributions (*P*_*count*_, *n*_*i*_, and *N*_obs_) that parameterize the simulations. Empirical discovery curves are computed for the full training set as well as all annotated lines, while the fitted model is used to extrapolate the discovery curve to the complete split-GAL4 collection (|*L*| = 3060). The empirical discovery curve for the subset is shown here for visualization purposes only, as the subset serves exclusively to compute *P*_count_, *n*_*i*_, *N*_obs_. All discovery curves represent the mean over Monte Carlo samples: for empirical data, curves are averaged over random orderings of underlying lines; for simulated data, they are averaged across independent simulation runs. **B** Model performance is evaluated by how well the discovery curve is extrapolated beyond the lines used for fitting. The prediction error is computed at |*L*_ann_| (all annotated lines, including those held out from fitting) and estimated at |*L*| (all lines).

#### Extrapolation

To improve the robustness of the simulation with respect to the random partitioning of the annotated split-GAL4 lines, we employed Monte Carlo cross-validation. Specifically, we fitted *M* = 100 models using independently sampled training and validation sets. For each model *m* ∈ {1, …, *M*}, we computed 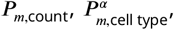, and 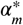 as described above. For each fitted model, we generated *R*_sim_ = 100 simulated discovery curves 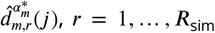, for the complete collection of lines |*L*| Because Monte Carlo cross-validation repeatedly resamples the annotated split-GAL4 lines, each annotated line contributes to multiple training and validation sets across the *M* fitted models. To leverage all annotated lines and improve the robustness of the final estimates, we averaged all *M* × *R*_sim_ = 10,000 simulated discovery curves across the fitted models. The predicted extrapolated number of unique cell types was then obtained from the averaged simulated discovery curve at *j* = |*L*|, and the corresponding coverage estimate was calculated by dividing this value by the total number of known cell types (|*C* | ).

Region-wise analyses were performed analogously, but were restricted to annotations and cell types localized to individual anatomical compartments. Supplemental Files [2] and [3] report the mean coverage estimates for the brain, the VNC, and their corresponding anatomical regions (cf. columns *num cell types simulated* and *coverage from simulated*).

#### Validation

Each of the *M* = 100 fitted models was evaluated for how well it extrapolates the discovery curve beyond its training lines. To this end, we compare the mean empirical discovery curve 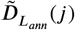 for all annotated lines with its model-specific mean simulated discovery curve 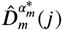 at position *j* = |*L* _*ann*_|, which corresponds to an extrapolation to ∼33% more lines, namely the previously held-out ones. We refer to this difference as the prediction error:

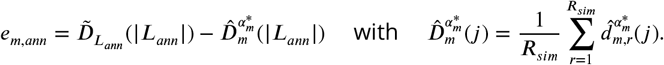

To extrapolate the error to the full line collection, we first compute the standardized distance *z* of *e*_*m,ann*_ with respect to the distribution of all simulated discovery curves of a model *m* at position *j* = |*L*_*ann*_|. Let *σ*_*m*_(*j*) denote the standard deviation of the *R*_sim_ simulated discovery curves for fitted model *m* at position *j*. For each model, we therefore computed

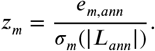

We now assumed that the same standardized distance *z* applies at both positions, *j* = |*L*_*ann*_| and *j* = |*L*| = 3060. Thus the prediction error for the complete split-GAL4 line collection was estimated by applying it at |*L*|:

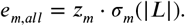

The final reported error was then obtained as mean absolute error (MAE) across all models:

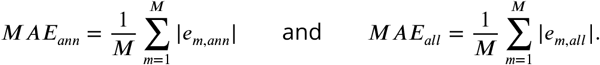

Figure 8**B** illustrates the prediction error *e*_*m,ann*_ and *e*_*m,all*_. Supplemental Files [2] and [3] report the mean absolute errors for both the annotated set and the full split-GAL4 line collection (cf. columns *Mean absolute error for all annotated lines* and *Mean absolute error for all lines*).

## Supporting information

Supplemental File 1: Updated cell type annotations for split-GAL4 collection

Supplemental File 2: Annotations and coverage analysis for brain

Supplemental File 3: Annotations and coverage analysis for VNC

Supplemental File 4: Sub cell type groupings

Supplemental File 5: Annotation guidelines and visualization example

Supplemental File 6: Blinded proofreading testing

Supplemental File 7: Compare new to published annotations

## Acknowledgments

We thank Jody Clements, Cristian Goina, Hideo Otsuna, Konrad Rokicki, and Robert Svirskas for the addition of cell type information to NeuronBridge, along with their ongoing work to add new LM & EM data. We thank Aljoscha Nern for feedback on the manuscript. We also thank Claudia Simone Winklmayr and Andreas Mardt for helpful discussions during the conceptualization of the statistical model. We thank the Janelia FlyLight Project Team and collaborators for creating the analyzed fly line and image collection. We thank the Janelia FlyEM Project Team and collaborators for creating the analyzed Hemibrain and MANC connectomes. This work was supported in part by the Janelia Visiting Scientist Program, and by the Howard Hughes Medical Institute. This article is subject to HHMI’s Open Access to Publications policy. HHMI lab heads and project team leads have previously granted a nonexclusive CC BY 4.0 license to the public and a sublicensable license to HHMI in their research articles. Pursuant to those licenses, the author-accepted manuscript of this article can be made freely available under a CC BY 4.0 license immediately upon publication.

## Supplemental Files

- Supplemental File 1: Updated cell type annotations for split-GAL4 collection
- Supplemental File 2: Annotations and coverage analysis for brain
- Supplemental File 3: Annotations and coverage analysis for VNC
- Supplemental File 4: Sub cell type groupings
- Supplemental File 5: Annotation guidelines and visualization example
- Supplemental File 6: Blinded proofreading testing
- Supplemental File 7: Compare new to published annotations

### Algorithm 1

Simulation of Cell Type Discovery

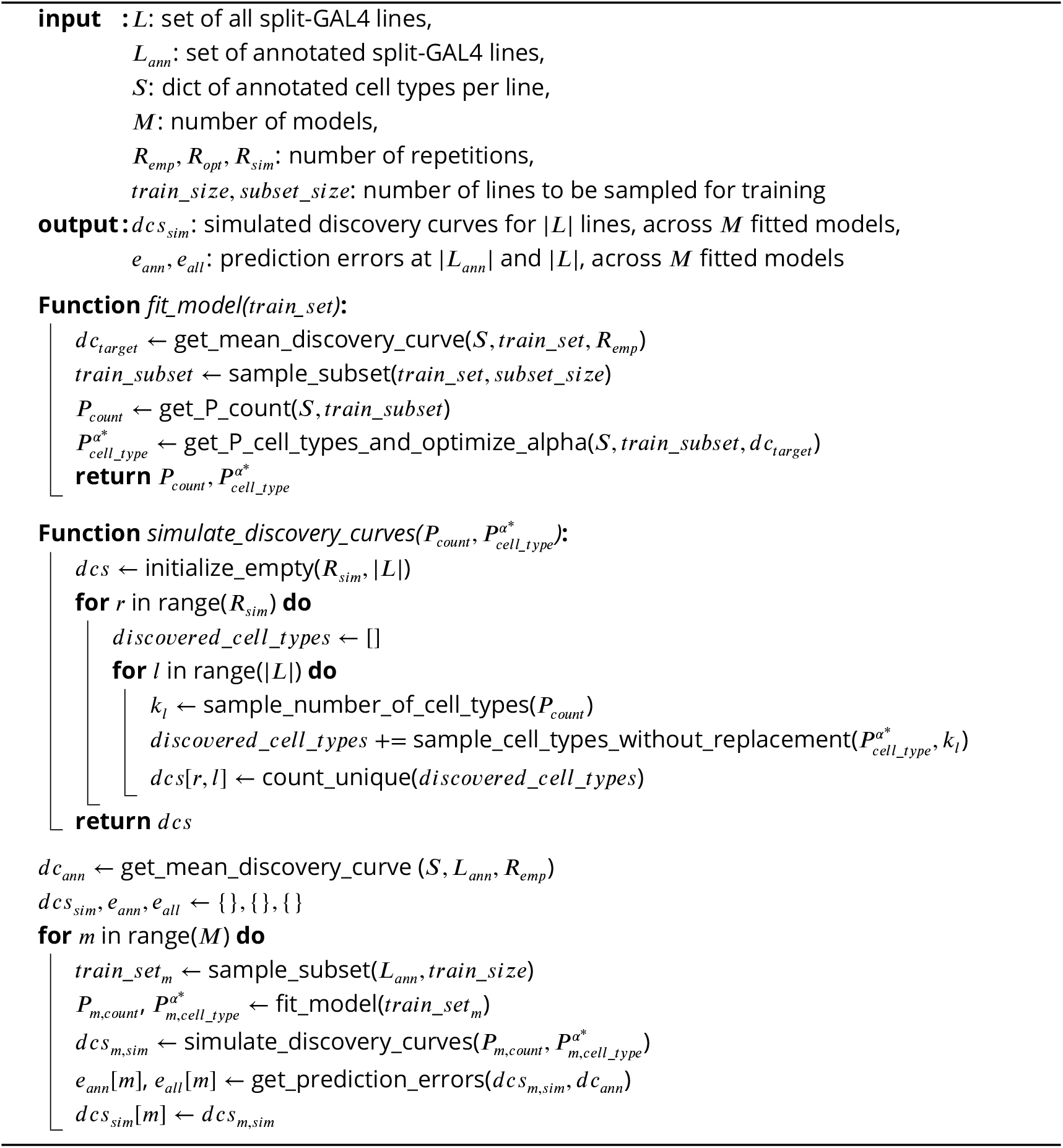

## Data availability

The code for PatchPerPixMatch is available at https://github.com/Kainmueller-Lab/PatchPerPixMatch, and the code for the coverage analysis is available at https://doi.org/10.25378/janelia.34535850. While Supplemental Files [2] and [3] provide the consolidated *Confident* and *Candidate* cell type-line matches used in this study, the underlying manually proofread automated search results, i.e., the *Confident, Candidate*, or *False* label assigned to each EM body ID-to-line match, are available in the coverage analysis repository.

## Author contributions

Lisa Mais: conceptualization, methodology, software, validation, formal analysis, investigation, data curation, writing—original draft, visualization. Claire Managan: methodology, validation, investigation, data curation. Brianna Yarbrough: investigation. Gary Patrick Hopkins: investigation. Jennifer Rivas Salinas: investigation. Shirley A. Lauchie: investigation. Peter Hirsch: visualization, writing-review. Christoph Karg: methodology, formal analysis, supervision. Stephan Saalfeld: conceptualization. Wyatt Korff: conceptualization, supervision, funding acquisition. Gudrun Ihrke: methodology, supervision. Dagmar Kainmueller: conceptualization, methodology, supervision, funding acquisition, writing-review. Geoffrey W Meissner: conceptualization, methodology, validation, data curation, writing—original draft, project administration.

