## Supplemental File 5: Annotation guidelines and visualization example for "Neuron types and distribution of a Drosophila driver line collection"

### Workflow

- Select a line to work on
- If a line is empty (no labeled neurons) or only has expression in the optic lobes, skip
- If a line appears to be too difficult, flag for senior annotator

### Possible annotations - definitions and usage

| Confident | The result is a match for a cell type or neuron that appears in the line.  There should only be one cell type annotated as TRUE for each cell type in the line. |
| --- | --- |
| FALSE | The result is NOT a match for a cell type or neuron that appears in the line. |
| Candidate | Cannot know for sure whether the result is a match for a cell type or neuron that appears in the line.  If there are multiple cell types or subtypes that could be true hits, label them all as “Candidate" |

### PDF examples

#### PDF overview

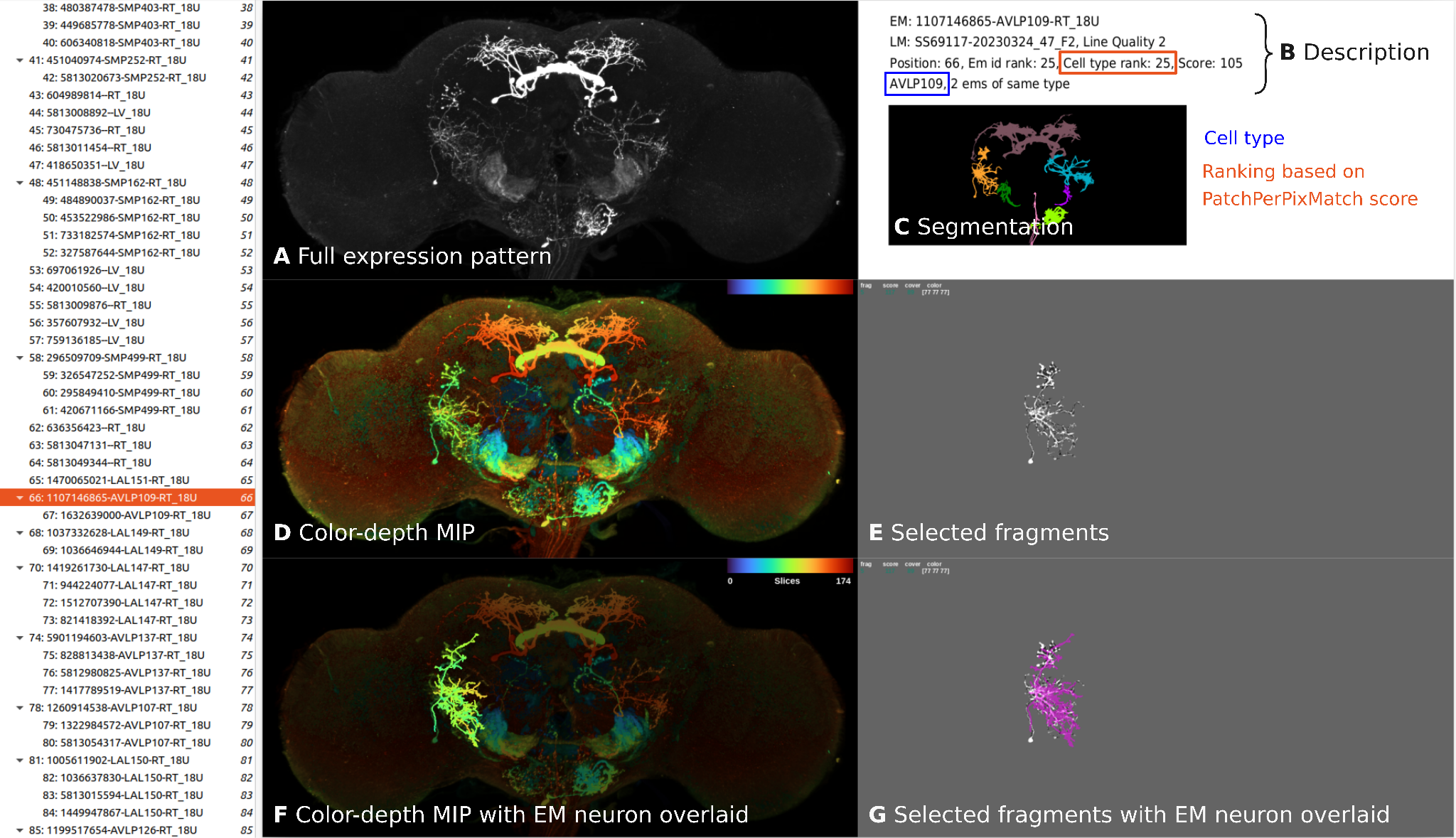

Visualization of PatchPerPixMatch search results for one split-GAL4 line:

**A** Maximum intensity projection (MIP) of the full expression pattern of the split-GAL4 sample.

**B** Description of the matching candidate, including EM body id, LM sample name, cell type rank across all matched EM cell types, as well as expert annotations if existent.

**C** MIP of instance segmentation masks where each color represents one individual component.

**D** Color-depth MIP (i.e. projection image with depth encoded as color) of the full expression pattern.

**E** Full expression pattern MIP masked by best-matching PatchPerPix fragments.

**F** Color-depth MIP of full expression pattern, dimmed and overlaid with color-depth projection of matched EM neuron morphology.

**G** Full expression pattern MIP masked by best-matching PatchPerPix fragments (pruned if the match stems from a pruned run), overlaid with EM neuron morphology in magenta.

#### Visualization of published annotations

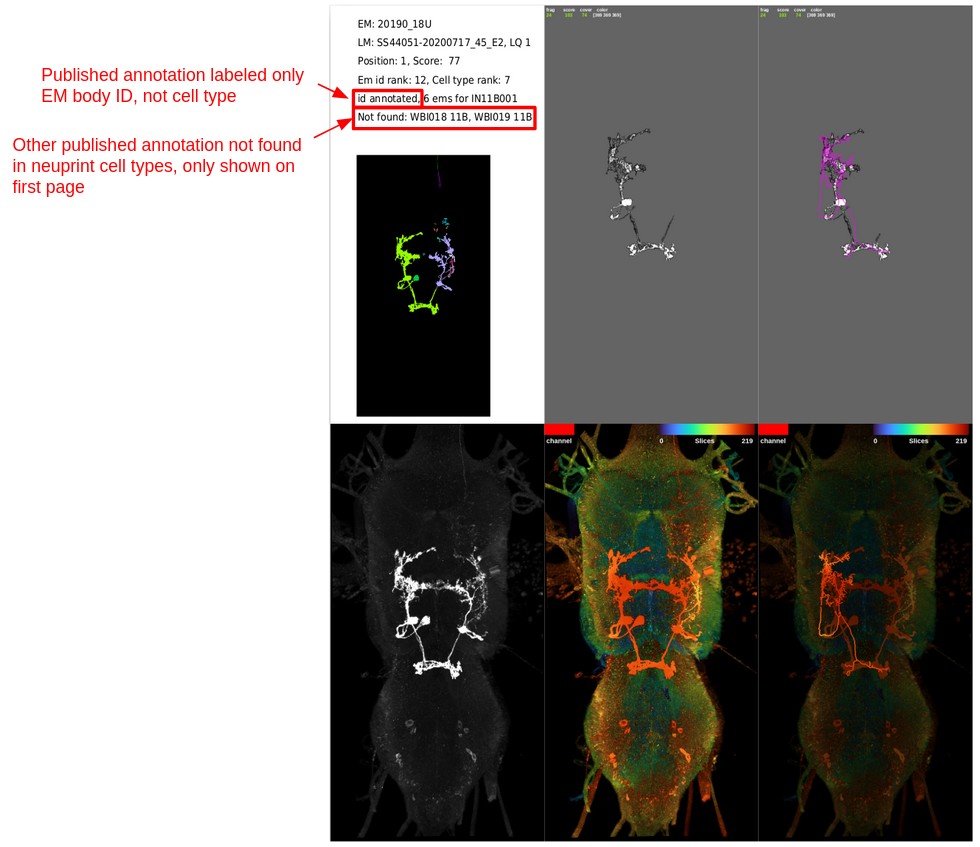
